# Patterns, predictors, and consequences of dominance in hybrids

**DOI:** 10.1101/818658

**Authors:** Ken A. Thompson, Mackenzie Urquhart-Cronish, Kenneth D. Whitney, Loren H. Rieseberg, Dolph Schluter

## Abstract

Are first-generation (F_1_) hybrids typically intermediate for all traits that differentiate their parents? Or are they similar to one parent for most traits, or even mismatched for divergent traits? Although the phenotype of otherwise viable and fertile hybrids determines their fate, little is known about the general patterns, predictors, and consequences of phenotype expression in hybrids. To address this empirical gap, we compiled data from nearly 200 studies where traits were measured in a common environment for two parent populations and F_1_ hybrids. We find that individual traits are typically halfway between the parental midpoint and one parental value (i.e., hybrid trait values are typically 0.25 or 0.75 if parents’ values are 0 & 1). When considering pairs of traits together, a hybrid’s multivariate phenotype tends to resemble one parent (pairwise parent-bias) about 50 % more than the other while also exhibiting a similar magnitude of trait mismatch due to different traits having dominance in conflicting directions. We detect no phylogenetic signal nor an effect of parental genetic distance on dominance or mismatch. Using data from an experimental field planting of recombinant hybrid sunflowers—where there is among-individual variation in dominance and mismatch due to segregation of divergent alleles—we illustrate that pairwise parent-bias improves fitness while mismatch reduces fitness. Importantly, the effect of mismatch on fitness was stronger than that of pairwise parent-bias. In sum, our study has three major conclusions. First, hybrids between ecologically divergent natural populations are typically not phenotypically intermediate but rather exhibit substantial mismatch while also resembling one parent more than the other. Second, dominance and mismatch are likely determined by population-specific processes rather than general rules. Finally, selection against hybrids likely results from both selection against somewhat intermediate phenotypes and against mismatched trait combinations.

## Introduction

When divergent populations occur in sympatry, they might mate and form hybrids (Mallet 2005). If those hybrids are viable and fertile, whether they survive and reproduce depends on their ability to persist under prevailing ecological conditions. Because selection against hybrids limits gene flow between parents (Harrison 1993), understanding the mechanisms underlying hybrid performance under ecologically-relevant conditions is key to understanding post-zygotic isolation (Barton and Hewitt 1985; Gompert et al. 2017). Quantifying general patterns of phenotype expression in hybrids is of interest because such patterns can shed light on the mechanisms underlying selection against hybrids. For example, if hybrids resemble one parent they could thrive in that parent’s niche and readily back-cross (Mallet 1986). Alternatively, if hybrids are phenotypically intermediate for all traits, or possess mismatched trait combinations due to dominance in opposing directions, they might be unable to survive and reproduce in the available niche space (Arnegard et al. 2014; Cooper et al. 2018; Hatfield and Schluter 1999; Matsubayashi et al. 2010). Currently, little is known about general patterns of trait expression in hybrids.

Previous synthetic studies investigating hybrid phenotypes have mixed conclusions. Some authors suggest that hybrid intermediacy is the rule (Hubbs 1940) whereas others find that hybrids are better described as mosaics of parental and intermediate characters (Rieseberg and Ellstrand 1993). Such previous studies typically lacked a quantitative framework and/or focused on a single taxon (e.g., fish or plants), limiting our ability to arrive at general conclusions. In addition, previous studies of hybrid phenotype expression tend to use mostly data from domesticated taxa, wherein dominance is expected to be elevated compared to natural populations (Crnokrak and Roff 1995; Fisher 1931). Here, we use a geometric approach to quantify patterns of hybrid phenotypes in a way that is comparable across studies. By quantifying the ‘parent-bias’ across each pair of traits we determine the extent to which hybrids are intermediate or tend to resemble one parent more than the other. And by quantifying the ‘mismatch’ (also termed ‘opposing dominance’ [Matsubayashi et al. 2010; Nosil 2012]) we can determine the extent to which hybrids have mismatched combinations of divergent parental traits (i.e., resemble parent 1 for trait *x* but parent 2 for trait *y*).

In this article, we systematically document patterns of phenotype expression in hybrids, investigate the possible predictors of these patterns, and use experimental data to explore the fitness consequences of trait interactions in the field. We first summarize the results of a systematic literature review of nearly 200 studies that compared the phenotypes of hybrids and parents in a common environment. We then ask whether features of a cross—such as the genetic distance between, or taxon of, the parents—are associated with dominance. We then use data from an experimental planting of recombinant hybrid sunflowers to evaluate whether patterns of pairwise parent-bias and mismatch predict fitness in hypothesized directions. Our results provide insight into the mechanisms that might commonly underlie selection against hybrids in nature.

## Methods

In this section, we provide a brief summary of, and rationale for, our methodology. A detailed explanation of all methods, including a summary of the data sources, is given in the supplementary methods.

### Systematic review of dominance patterns in F_1_ hybrids

We conducted a systematic literature search and identified 198 studies from which we could collect data of at least one phenotypic trait measured in two parent taxa and their F_1_ hybrids in a common environment. We included studies that conducted crosses between wild-collected parental populations or laboratory populations with fewer than ten generations of captivity. Data from wild hybrids (i.e., not from controlled crosses) were only included if hybrids were genotyped. We aimed to include only traits with environment-dependent effects on fitness—traits plausibly under divergent, rather than directional, selection (sometimes called ‘non-fitness’ traits [Merilä and Sheldon 1999] or ‘ordinary’ traits [Orr and Betancourt 2001]). For example, traits such as ‘embryo viability’ are almost certainly under directional selection and were not included in our database. We excluded likely fitness components because developmental difficulties resulting from hybrid incompatibilities, or heterosis resulting from outbreeding, often affects such traits in hybrids (Coyne and Orr 2004). This choice to exclude fitness traits likely renders our analysis on dominance more conservative, since hybrid breakdown or heterosis would manifest as a transgressive phenotype. By contrast, traits such as ‘limb length’ might have particular values best suited to some environments and genetic backgrounds—it is implausible that such traits would always be selected to a maximum or minimum value. Data from back-cross (BC_1_ only) and F_2_ hybrids were collected when available, but primarily used to address a different question (see Thompson 2019).

The studies in our analysis spanned a range of taxa but included mostly vascular plants (34 %), arthropods (29 %), and vertebrates (30 %). We do not use formal phylogenetic comparative methods for our main inferences because we are not testing or proposing a causal model (Uyeda et al. 2018), and we generally do not use formal meta-analysis tools because we are not synthesizing experimental data.

Because our interest is specifically in quantifying patterns of dominance for traits that differentiate species, we first restricted our dataset to putatively divergent traits. We did this by retaining traits for which parents had statistically divergent phenotypes (*t*-test *P* < 0.05). In addition, we retained a small number of traits for which the parents were greater than 1 SD apart but not statistically distinguishable (using the smaller parental SD; see Fig. S1). After filtering traits, we converted all trait data that were published with a transformation (i.e., ln, square-root) to their original measurement scale because expectations are not the same on a log or squareroot scale as for raw units. We then put all traits in all studies on a common scale where one (arbitrarily determined) parent had a value of 0 for all traits and the other had a value of 1 (see Fig. 1). Because we do not make any assumptions about which trait value is ancestral or derived, dominance and recessivity of traits is indistinguishable. For example, a trait’s degree of dominance is the same whether the hybrid trait value is 0.2 or 0.8. Under an expectation of additivity, a hybrid would have a trait value of 0.5 for all traits. Importantly, however, if a hybrid had two traits with values [0.2, 0.8], the arithmetic mean phenotype is indeed 0.5 but the hybrid is not intermediate but rather mismatched. The hybrid in this case resembles one parent for the first trait and resembles the other parent for the second trait. This failure of simple averaging highlights the need dominance metrics based on the geometry of phenotype space.

**Fig. 1.**
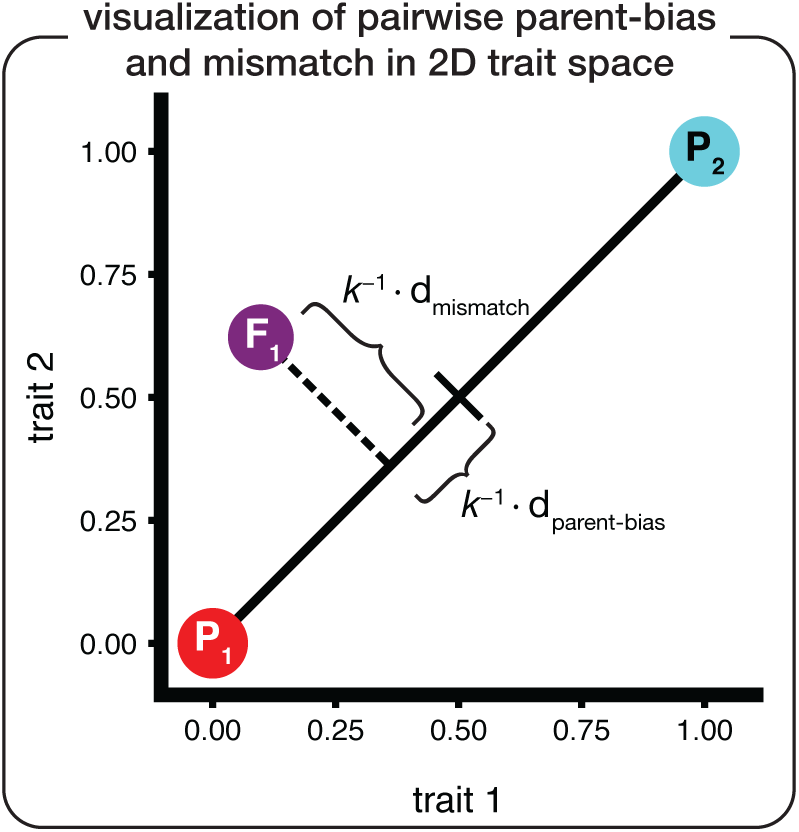
Visual overview of how 2-dimensional dominance metrics were calculated. When studies contained two or more divergent traits we calculated pairwise parent-bias (d_parent-bias_) and mismatch (d_mismatch_) of the hybrid (here F_1_ but could in principle be any hybrid generation) phenotype with respect to the line connecting the two parent phenotypes (P_1_ & P_2_). This procedure was repeated for every pair of traits. *k* is a scaling factor that renders the maximum value observed without transgression (i.e., d_mismatch_ when F_1_ trait values are [0, 1]; or pairwise d_parent-bias_ when F_1_ trait values are [0, 0]) equal to 1. For two traits, 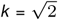 Values > 1 can result when traits are transgressive.

### Quantifying dominance in F_1_ hybrids

We quantified three types of dominance in the data (see Fig. 1). Within a cross, each dominance metric was scaled such that values of 0 indicate no dominance, values of 1 indicate the maximum dominance without transgressing the parental trait range, and values greater than 1 result from transgression.

The first dominance metric is ‘univariate’ dominance (d_univariate_), which considers traits individually. Specifically, d_univariate_ is the average deviation of trait values from 0.5 (the additive expectation) regardless of direction. For a single trait, this was calculated as:

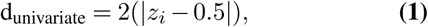

where *z*_*i*_ is the scaled mean phenotype of trait *i*. A d_univariate_ value of 0 results only when all traits have values of exactly 0.5. We calculated d_univariate_ for each trait within a cross and then took the mean of all traits within a cross.

The remaining two dominance metrics consider pairs of traits at a time and are calculated in 2-dimensions. We consider pairs of traits to increase the comparability of dominance values among crosses—dimensionality affects Euclidean distance metrics and therefore it is not possible to directly compare studies measuring different numbers of traits. For crosses where three or more divergent traits were measured, we calculated 2-dimensional dominance for each trait pair and then took the mean of all pairwise estimates as the value for that cross. For each pair of traits, we first determined the scalar projection, *b*, of the hybrid phenotype onto the line connecting parents (dashed line in Fig. 1). This projection is calculated as:

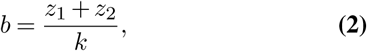

where *z*_1_ and *z*_2_ are the hybrid values for trait 1 and 2.

The second metric of dominance is pairwise parent-bias (d_parent-bias_), which captures deviation from bivariate intermediacy in the direction of either parent. This value is less than d_univariate_ when dominance values are variable among traits. For example, if a hybrid’s standardized phenotype is [0, 1] for traits 1 and 2, respectively, then the mean d_univariate_ = 1 but pairwise d_parent-bias_ = 0. Pairwise d_parent-bias_ has a minimum value of zero when dominance is equally strong in the direction of both parents and increases indefinitely as dominance increases in a manner that is biased toward one parent. For each pair of traits, pairwise parent-bias was calculated as:

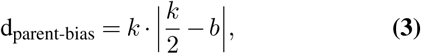

where *b* is the scalar projection from eqn. 2, *k* is a scaling factor 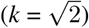 used to give a hybrid a phenotype with parental values for both traits a value of 1.

The final metric of dominance is pairwise mismatch, (d_mismatch_), which captures the perpendicular distance between the mean hybrid phenotype and the line connecting parental mean phenotypes. d_mismatch_ has a minimum value of zero when the hybrid phenotype is on the line connecting parents and increases indefinitely as the variance in dominance among traits increases. This metric captures what Rieseberg et al. (2003) describe as ‘mosaicism’, but on a continuous scale. For each pair of traits, mismatch was calculated as:

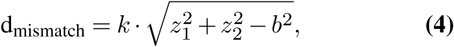

where, *z*_1_ and *z*_2_ are as in eqn. 1, and *b* and *k* are as in eqn. 3. Within each cross, both pairwise d_parent-bias_ and d_mismatch_ were calculated for every pair of traits and then averaged across all pairs.

### Testing possible predictors of dominance in F_1_ hybrids

We explored several possible predictors of dominance motivated by previous results and theoretical predictions. For example, previous studies have determined that genetic distance between the parents affects the frequency with which traits transgress the parent range (Stelkens and Seehausen 2009), a pattern that should be captured by our d_univariate_ metric. To determine if genetic distance affects dominance in our analysis, we computed genetic distance using gene sequence data and tested whether it was associated with any metric of dominance. Because genetic distance was calculable for less than one quarter of studies, we also compared dominance metrics between intraspecific and interspecific crosses—the underlying assumption being that genetic distance between parents is less in the former than in the latter.

If there exist fundamental differences in the genetics of adaptation among taxa, such differences could manifest as dominance. For example, self-fertilisation, which is more common in plants than animals, is expected to favour the fixation of recessive alleles (Charlesworth 1992), which could bias hybrid phenotypes away from intermediacy compared to outcrossing animals. Various taxon-specific reviews have arrived at different conclusions that suggest there might be differences between groups (Hubbs 1955; Rieseberg and Ellstrand 1993). To test whether there might be variation in dominance between taxa, we built a phylogeny encompassing nearly all studies in our dataset (Fig. S2) and tested for phylogenetic signal in dominance metrics. We also tested whether there are differences in dominance between predefined taxonomic groups such as plants and animals.

We finally tested specific hypotheses about phenotypic variation and parent-of-origin effects. If maternal effects are common, for example, then hybrid trait values might tend to resemble the maternal parent more than the paternal—this is testable in the present dataset because many crosses (*n* = 96) were conducted in both directions. We also investigated whether hybridization increases phenotypic variance. Variance in recombinant hybrids (BC_1_s and F_2_s) is expected to exceed that of parents. By contrast, phenotypic variance in F_1_ hybrids is expected to equal that of parents unless there are interactions between divergent parental alleles that reduce developmental stability in hybrids (Hochwender and Fritz 1999).

### Fitness consequences of parent-bias and mismatch in recombinant sunflowers

The above analyses were motivated by the hypothesis that, compared to a hybrid that is a perfect intermediate, hybrids resembling parents should fare well and hybrids that are mismatched should fare poorly. However, there is no way to test the fitness consequences of parent-bias and mismatch in the data synthesized from the literature because no studies have both individual-level phenotype and fitness data in the field. The best way to investigate the fitness effects of parent-bias and mismatch is to examine a population of recombinant hybrids—wherein individuals exhibit quantitative variation in the degree of parent-bias, mismatch, and fitness—and then to use these resulting data to test whether dominance metrics are associated with fitness.

To evaluate the fitness effects of parent-bias and mismatch, we leveraged data from a field experiment in annual sunflowers (*Helianthus*). Briefly, 475 *H. a. annuus* × *H. debilis* BC_1_ hybrids were planted alongside individuals of both parental species in central Texas. Fitness (seed number) as well as 30 architectural, floral, ecophysiological, phenological, and herbivore resistance traits were measured. We applied the same trait selection and filtering criteria as in the systematic review and retained 20 traits (see Table S1 for trait details). The data from this experiment have been previously published (Whitney et al. 2006, 2010).

For each plant, we calculated pairwise d_parent-bias_ and d_mismatch_ and then took the average across all trait pairs. pairwise d_parent-bias_ and d_mismatch_ (*r* = 0.762, *P* < 0.001; Fig. S3) because many traits in this dataset are transgressive and high single-trait dominance is the cause of both parent-bias and mismatch. Therefore, we investigated their respective effects on fitness using multiple linear regressions of the form:

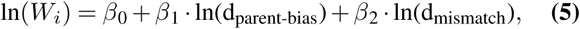

where in this case d_parent-bias_ and d_mismatch_ are the mean individual values averaged across each trait pair. We also ran the same multiple regression for each pair of traits separately, and asked whether the sign of regression coefficients (*β*_1_ and *β*_2_) were consistent with those observed in the analysis of mean pairwise parent-bias and mismatch.

## Results

### Patterns of dominance in F_1_ hybrids

Among unique crosses in the dataset, the mean univariate dominance (d_univariate_ ± 1 SE) measured in F_1_ hybrids was 0.791 ± 0.0776 (Fig. 2A; median = 0.549), which suggests that the average trait is not intermediate but rather more than halfway between intermediate and parental. In 20 % of crosses, the mean d_univariate_ was > 1, indicating transgression. The mean pairwise parent-bias (d_parent-bias_ among crosses was 0.679 ± 0.010 (Fig. 2B; median = 0.445), and the mean pairwise mismatch (d_mismatch_) was 0.597 ± 0.101 (Fig. 2C; median = 0.311).

**Fig. 2.**
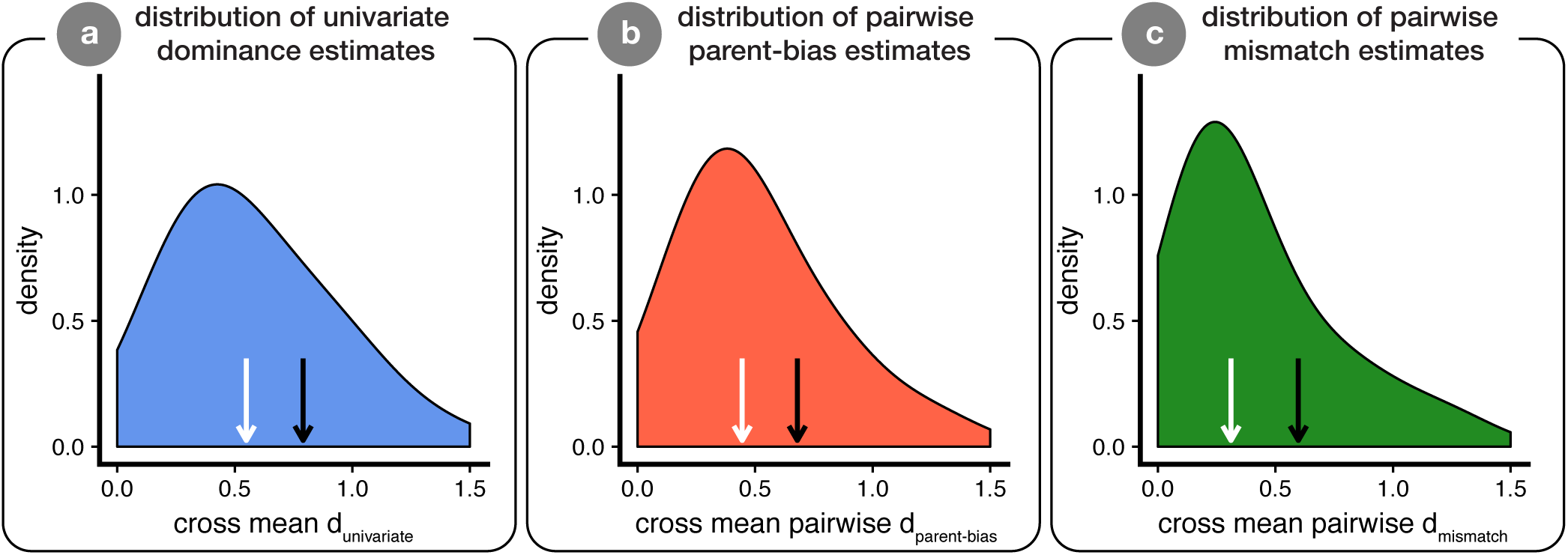
Patterns of dominance in F_1_ hybrids. The density plots (*y*-axis standardized across panels) show the three main dominance metrics contained herein, with each cross contributing at most a single value per panel. Values of 0 indicate no dominance, values of 1 indicate the maximum without transgression, and values > 1 reflect transgression. The *x*-axis is truncated at 1.5, but the means (black arrows) and medians (white arrows) are calculated from the whole dataset (see Fig. S5 for the same figures with unconstrained axes and Fig. S4 for a summary of patterns when each cross contributes a median rather than mean value). Panel **a** shows the univariate dominance (d_univariate_; eqn. 1), panel **b** shows parent-bias (pairwise d_parent-bias_; eqn. 3), and panel **c** shows mismatch (pairwise d_mismatch_; eqn. 4). Panel **a** contains one value from all crosses (*n* = 233) while panels (**b**) and (**c**) only contain information from crosses wherein two or more traits were measured (*n* = 165). Density plots were generated using the `geom_density` function in `ggplot2` with twice the default smoothing bandwidth.

The above metrics of dominance are likely influenced by sampling error. This is because sampling error around an intermediate phenotype would appear as dominance when taking the absolute value of the difference between the observed mean phenotype and the mid-parent value. To determine the magnitude of dominance estimates caused by sampling error alone, we generated 1000 simulated datasets with identical structure, sample sizes, and standard deviations as the real data but where the true mean of F_1_ hybrid traits was exactly intermediate between the parents. We then calculated the median of the three dominance metrics for each simulated dataset. Across all simulations, the median d_univariate_ was 0.200 (0.349 units less than observed), the median pairwise d_parent-bias_ was 0.113 (0.332 units less than observed), and the median pairwise d_mismatch_ was 0.112 (0.200 units less than observed). No simulation run ever returned a median dominance metric as large as that observed in the real data (see Fig. S6). The simulation results indicate that the majority of our signal is biological rather than caused by sampling error.

### Predictors of dominance in F_1_ hybrids

We next investigated whether dominance patterns in F_1_ hybrids are associated with genetic distance and phylogeny. We found no significant associations between any metric of dominance and any metric of genetic distance (see detailed results in Figs. S7–S9). In addition, there was no evidence for phylogenetic signal in any dominance metrics (all *λ* < 0.00001, all *P* = 1), and no difference in any dominance metrics in comparisons of major clades (Fig. S9).

Because many crosses were conducted reciprocally (i.e., 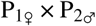 and 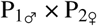), we could evaluate whether hybrids tend to resemble their maternal parent more than their paternal parent (or vice-versa). We found that 25.6 % of traits differed significantly (at *P* = 0.05) between cross directions. The mean magnitude of phenotypic difference between cross directions was 0.65 SDs (units of smaller parental SD). Within each cross that was conducted in two directions, we calculated the fraction of traits that exhibited maternal bias. We then used a one-sample *t*-test to test whether this fraction deviated significantly from 0.5. We found that traits of F_1_ hybrids tend to resemble the maternal parent about 57 % of the time. (*t*_94_ = 2.034, *P* = 0.0447, 95 % CI = [0.502, 0.657]).

Various authors have speculated that dominance tends to be in the direction of the higher trait value (e.g., Fisher 1931). We found that dominance was biased toward the higher value in 51.8 % of traits, which was not significantly different from the random expectation of 50 % (binomial test; *P* = 0.265).

Last, we were interested in determining whether hybrids had comparable or greater phenotypic variance compared to parents. If F_1_ hybrids typically suffer difficulties during development, they might have increased phenotypic variation relative to parents. We found that phenotypic variation in F_1_ hybrids was statistically indistinguishable from the average of the parental SDs (*P* = 0.271) (Fig. S11). F_2_ hybrids had, on average, 2.3× the phenotypic SD of the mean-parent SD (*P* < 0.0001) and first-generation back-crosses had 1.59× the variance of parents (*P* = 0.0001).

### Fitness effects of parent-bias and mismatch in recombinant sunflowers

We used data from an experimental planting of recombinant hybrid sunflowers to evaluate whether our metrics of pairwise parent-bias and mismatch explain variation in fitness in the field. We first asked if the mean pairwise parent-bias and mismatch were associated with fitness (eqn. 5). We found that parent-bias was positively associated with seed count (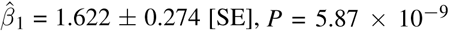; Fig. 3A), while mismatch had a negative association (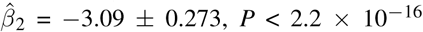; Fig. 3B). We detected no significant interaction term (*P* > 0.8) in a separate multiple regression model that tested for it. In this dataset, the fitness consequences of a unit change in d_mismatch_ were larger than the fitness consequences of an equivalent unit change in pairwise d_parent-bias_ (test of hypothesis that 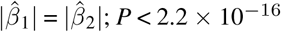). We note that pairwise trait-correlations were typically quite low in these data (mean | *ρ*| = 0.149; Fig. S10). Together, mean pairwise parent-bias and mismatch explained 27.7 % of the total variation in seed count. The fitness consequences of pairwise mismatch were greater than selection on each of the 20 traits individually, and the fitness consequences of pairwise parent-bias were greater than 19 of 20 traits.

**Fig. 3.**
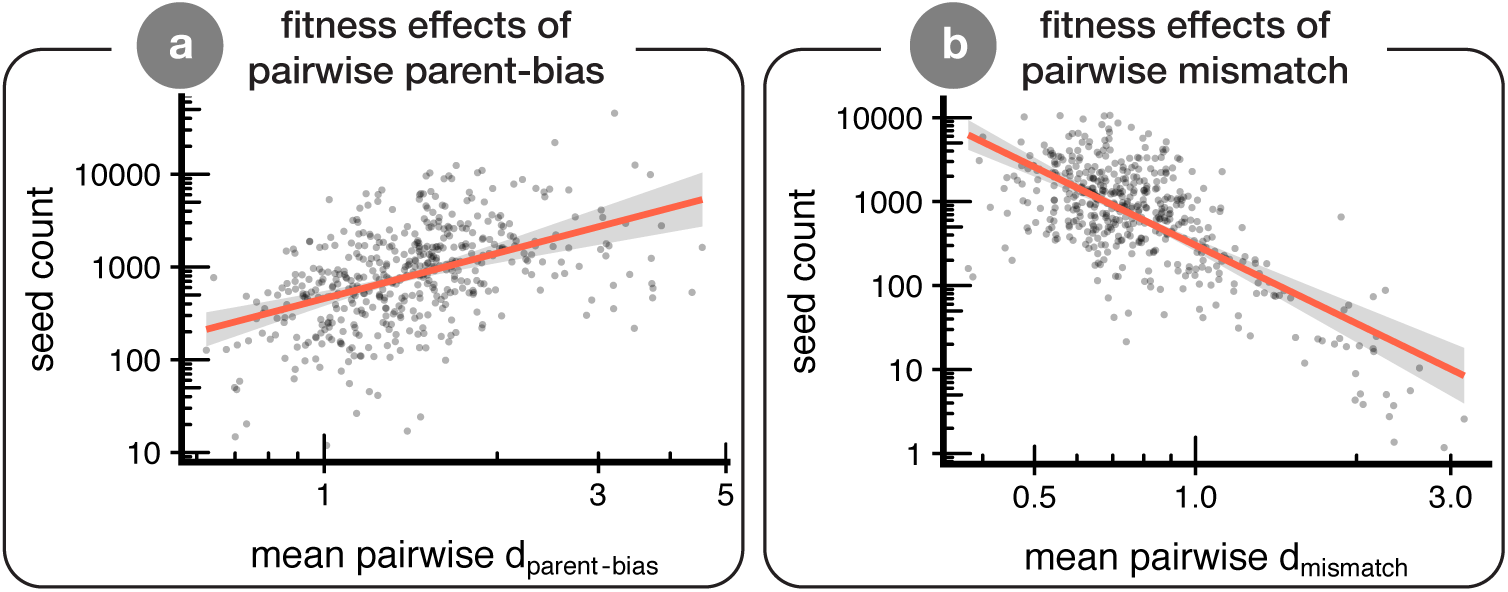
Effect of parent-bias and mismatch on fitness in *H. a. annuus* × *H. debilis* BC_1_ hybrid sunflowers growing in the field. The points are partial residuals extracted from a multiple regression using `visreg`. Each point represents one individual hybrid plant (*n* = 475). Both axes are log_10_ transformed. Panel **a** illustrates the effect of dominance via parent-bias and panel **b** illustrates the effect of dominance via mismatch.

We also evaluated dominance-fitness relationships for each pair of traits separately. This analysis is heuristic because pairs of traits are not independent, but we present it to complement the above results. Considering only statistically significant coefficients, pairwise d_parent-bias_ improved fitness for 62.3 % of trait pairs and d_mismatch_ reduced fitness for 84 % of trait pairs (Fig. S12). Thus, the fitness and pairwise d_parent-bias_ and d_mismatch_ is consistent between analyses of an individual’s mean value averaged over all trait pairs and when considering trait pairs individually.

## Discussion

In this article, we compiled data from studies that measured phenotypic traits in F_1_ hybrids to characterise general patterns of hybrid trait expression. We then investigated whether the observed dominance could be predicted by several variables including genetic distance and phylogeny. Last, we tested whether parent-bias and mismatch were associated with fitness in the field. From the data, it is clear that dominance is common. This is most clearly demonstrated by our finding that, in F_1_ hybrids, individual traits are, on average, halfway between the parental midpoint and one parent’s phenotype. In addition, there was no association between dominance and any predictor variable, suggesting that it will be difficult to make accurate predictions about the patterns of dominance for any individual cross. In the sunflower data, both pairwise parent-bias and mismatch affected fitness in recombinant hybrid sunflowers in the predicted direction. We discuss these results in the context of previous research on dominance and trait expression in hybrids, and highlight the implications for speciation research.

### Genetic underpinnings of dominance and mismatch

Why is dominance so commonly observed in F_1_ hybrids? Unfortunately, because we do not know which trait values are derived vs. ancestral, most theories on the evolution of dominance (e.g., Haldane’s sieve [Haldane (1924, 1927)]) cannot be applied. In any case, inter-population phenotypic divergence is likely underpinned by many quantitative trait loci (QTL) and our results hint at two general features of such QTL. First, the alleles used for adaptation do not exhibit strict additivity. Supporting this, Miller et al. (2014) quantified dominance of QTL underlying marine-freshwater phenotypic divergence in threespine stickleback (*Gasterosteus aculeatus*) and found that the majority of QTL had some dominance effects—although there was no directional bias toward either derived or ancestral conditions, likely because adaptation was from standing variation rather than *de novo* mutation (Orr and Betancourt (2001)). And second, different traits have imbalanced dominance coefficients, such that some traits have QTL dominance effects favouring one parent while other traits have effects biased toward the other. Such patterns would result most readily when diverging populations experience selection on different traits in their respective populations—that is, when adaptive divergence proceeds in orthogonal directions in phenotype space.

### Patterns and predictors of dominance

Our results corroborate some previous findings but are inconsistent with others. Hubbs (1940) suggested that fishes show additive inheritance “as a very general rule” whereas Rieseberg and Ellstrand (1993) suggested plant hybrids are best characterised as being “a mosaic of both parental and intermediate morphological characters rather than just intermediate ones”. Our quantitative analysis paints a picture more akin to mosaicism than strict intermediacy. In addition, we find no evidence for any major differences in dominance among taxonomic groups, which suggests the choice of study taxon does not bias estimates of dominance.

Stelkens and Seehausen (2009) found that the genetic distance between parents in interspecific crosses was positively correlated with transgression frequency—the tendency for traits to fall outside the range of parental values. We used an almost entirely independent dataset and did not find such a pattern. Perhaps the most likely cause of this discrepancy is that Stelkens and Seehausen (2009) considered traits reflecting ‘intrinsic’ hybrid incompatibility (e.g., small body size due to poor condition or low seed production due to inviable ovules) and F_1_ hybrid heterosis (e.g., larger body size or high seed count due to overcoming inbreeding depression in parents). Incompatibility increases with parental genetic divergence Matute et al. 2010; Moyle and Nakazato 2010; Orr 1995; Wang et al. 2015), and heterosis seems to as well until inviability becomes substantial (Wei and Zhang 2018). Importantly, such inviability and heterosis would manifest as transgression using our approach. Because we only consider wild outbred taxa and traits that are putatively under divergent selection, and other studies considered mostly fitness components in domesticated and lab-adapted populations, the mechanisms linking genetic distance with transgression in earlier studies do not apply to the present dataset.

### Fitness consequences of mismatch

Our results clarify the potential for dominance to have a role in driving progress toward speciation. Our findings challenge the conjecture that reduced F_1_ fitness is due only to phenotypic intermediacy and hybrids ‘falling between parental niches’. This is simply because F_1_ hybrid trait values are typically not the mid-parent value. Rather, F_1_ hybrids often possess novel multivariate phenotypes that are best viewed as being more similar to one parent than the other, and also as being moderately mismatched. In nature, the phenotype of an organism is an integrated suite of traits that function together to influence performance and ultimately fitness (Arnold 1983; Brodie 1992). Because mismatch, caused by dominance in conflicting directions among traits, breaks up suites of integrated traits, this likely causes the resulting hybrids to be poorly suited to any environment.

In the sunflower data, we found that pairwise parent-bias improved fitness and mismatch reduced fitness. At present, it is not clear how general this finding is. Importantly, mismatch was more detrimental than parent-bias was beneficial. To the extent that mismatched combinations are selected against, and matched combinations are selected for, our results suggest that low F_1_ hybrid fitness in nature is caused both by partial-intermediacy and partial-mismatch. F_1_ hybrids are likely closer in phenotype to one parent than the other, and yet at the same time have some traits resembling the less-similar parent which might render them unable to survive and reproduce in the similar-parent’s niche. It would be valuable to conduct more field experiments with recombinant hybrids to arrive at generalities in the ways that parent-bias and mismatch affect fitness.

## Conclusion

In this study we synthesized data from nearly 200 studies and over 230 crosses and described general patterns of phenotype expression in F_1_ hybrids. Compared to previous studies with a similar goal, the distinguishing features of our analysis are that we used continuous trait data rather than categories, looked across several major clades, and examined divergently-selected traits in wild organisms. For individual traits, reasonably high dominance is the rule rather than the exception. Previous studies have documented the phenomenon where dominance acts in opposite directions for different traits (Matsubayashi et al. 2010). We built on these previous studies by quantifying mismatch using simple 2D geometry and demonstrating that such patterns affect most hybrids to a fairly substantial degree.

Previous authors have qualitatively drawn a link between trait mismatches and hybrid fitness (e.g., Arnegard et al. 2014; Cooper et al. 2018), and we built on these earlier results to link individual-level mismatch metrics to fitness. This result contributes to a growing literature on trait interactions in hybrids, and we suggest that future studies use our approach (or a complementary approach) to test the fitness consequences of mismatch directly. Such trait interactions are similar to Bateson-Dobzhansky-Muller hybrid incompatibilities (BDMIs) with fitness consequences mediated via ecology. Ecological BDMIs have the opportunity to affect many F_1_ hybrids and could be a major mechanism of extrinsic post-zygotic isolation. Only field observations and experiments can provide the data that are necessary to test this hypothesis.

## Acknowledgements

Feedback from S. Barrett, D. Irwin, M. Pennell, and S. Otto benefited the manuscript. K. Davis, N. Frasson, J. Heavyside, K. Nikiforuk, and M. Urquhart-Cronish assisted with the literature search (KD, NF, JH, M U-C) and/or data collection (KN & MU-C). We are grateful to dozens of authors who responded to our requests for data. The authors were supported by Genome British Columbia, Genome Canada, the Global Crop Diversity Trust, the United States National Science Foundation (NSF DEB 1257965 [to KDW & LHR]), NSERC (discovery grants to LHR & DS, graduate scholarships to KAT & MUC), the University of British Columbia (four-year fellowships and tuition funds to KAT & MUC), and the Izaak Walton Killam Memorial Fund for Advanced Studies (KAT). We are grateful to the organizers of the Hamburg Hybridization symposium for providing a fantastic venue for facilitating collaboration. R. Henriques created the bioR*X*iv LATEXtemplate.

## Author contributions

KAT concieved of the systematic review and designed the protocol with input from MUC and DS. KAT and MUC screened studies and collected data, and KAT contacted authors for data if necessary. KAT checked all data for accuracy. KDW had the idea to explore the fitness effects of parent-bias and mismatch in sunflowers, and KDW & LHR contributed sunflower data. KAT analysed the data and wrote the paper with input and contributions from all authors.

## Data accessibility

All data and analysis code used in this article will be deposited in a repository (e.g., Dryad) following publication.

## Supplementary material 1: Supplementary methods

### Search strategy

We searched the literature for studies that made measurements of traits in F1 hybrids and their parents. To identify studies for possible inclusion, we conducted a systematic literature search using Web of Science. We included all papers that resulted from a general topic search of “Castle-Wright”, and from a topic search of “F_1_ OR hybrid OR inherit*” in articles published (from any year) in *Evolution, Proceedings of the Royal Society B, Journal of Evolutionary Biology, Heredity*, or *Journal of Heredity*. These journals were selected because a preliminary search indicated that they contained nearly half of all suitable studies. These searches returned 82 of the 198 studies deemed suitable after screening. The literature search included all studies published until the end of 2017.

To be more comprehensive, we conducted additional systematic searches using the same keywords in articles citing influential and highly-cited publications (Bradshaw et al. 1998; Churchill and Doerge 1994; Coyne and Orr 2004; Dobzhansky 1937; Grant 1981; Hatfield and Schluter 1999; Hubbs 1955; Lande 1981; Lynch and Walsh 1998; Mayr 1963; Schluter 2000; Tave 1986). The full literature search results are available in the archived data. The initial search returned 14048 studies, and after removing duplicates this left 11,287 studies to be screened for possible inclusion.

### Comments on systematic nature of review

We attempted to follow PRISMA (Moher et al. 2009) guidelines to the best of our ability. Most of the criteria have been addressed above but a few other comments are warranted. In particular, we have no reason to suspect that any bias was introduced about dominance. This is because no studies seemed to have *a priori* hypotheses about such patterns. Accordingly, we do not believe that our estimates suffer from a file drawer problem. We emphasise that a formal meta-analytic framework—wherein data from multiple studies are aggregated with various weights—is not appropriate because we are not comparing studies that had any experimental treatment. Because the studies in our dataset do not have anything resembling an ‘effect size’, a simple summary across all of them is most appropriate.

### Evaluation of studies

We required studies to meet several criteria to merit inclusion in the database. First, the study organisms had to originate recently from a natural (i.e., ‘wild’) population. This is because dominance patterns in domestic species differ substantially from non-domesticated species (Crnokrak and Roff 1995) and because we were explicitly interested in documenting patterns as they occur in nature. We therefore excluded studies of crops or horticultural varieties, domestic animals, laboratory populations that were > 10 (sexual) generations removed from the wild, or where populations were subject to artificial selection in the lab. If populations were maintained in a lab for more than 10 generations but were found by comparison to still strongly resemble the source population, we included the study (*n* = 2 studies). We also excluded studies where the origin of the study populations was ambiguous. Hybrids had to be formed via the union of gametes from parental taxa, so we excluded studies that used techniques like somatic fusion. The ancestry of individual hybrids also had to be clear. This was always obvious in the case of laboratory crosses but was difficult in some studies of wild, naturally formed, hybrids. Many studies reported phenotypes of natural hybrids, for example in hybrid zones, and we did not include these studies unless the hybrid category (i.e., F_1_, F_2_, BC_1_) was confidently determined with molecular markers (typically over 95 % probability, unless the authors themselves used a different cut-off in which case we went with their cut-off) or knowledge that hybrids were sterile and thus could not be beyond the F_1_).

We are explicitly and exclusively interested only in divergently-selected traits. (A similar study of fitness components would be interesting but is beyond the scope of the present study.) By definition, these so-called ‘ordinary’ traits (Orr 2001) should be under stabilizing selection at their optimum, whereas traits that are direct fitness proxies are those that are likely under directional selection and have no optimum (Merilä and Sheldon 1999; Schluter et al. 1991). Thus, we required studies to report measurements of at least one ‘ordinary’ trait. In most cases it was possible to evaluate this distinction between fitness proxies and ordinary traits (hereafter simply ‘traits’) objectively because authors specifically referred to their reported traits as components of fitness, reproductive isolating barriers, or as being affected by (non-ecological; i.e., ‘intrinsic’) hybrid incompatibilities. In some cases, however, we made the distinction ourselves. If particular trait values could be interpreted as resulting in universally high fitness, for example resistance to herbivores or pathogens, this trait was not included. Said another way, if particular trait values could be seen as ‘good’ and others as ‘bad’, we did not include this trait. The majority of cases were not difficult to assess, but we have included reasons for excluding particular studies or traits in the database screening notes (see Data accessibility). Since fitness component traits are likely to exhibit heterosis or incompatibilities, a more inclusive approach would increase all of our metrics of dominance with the possible exception of pairwise d_parent-bias_.

Traits had to be measured in a quantitative manner to be included in the dataset. For example, if a trait was reported as being in categories related to parents or intermediacy (‘parent-like’ or ‘intermediate’), we did not include it. Some traits such as mate choice must often be scored discretely (in the absence of multiple trials per individual), even though the trait can vary on independent trials. Accordingly, we included discretely scored traits—like mate choice—when it was possible in principle to obtain a different outcome on independent trials. Such traits are recorded as 0s and 1s, but hybrids can be intermediate if both outcomes occurred with equal frequencies. We included traits where authors devised their own discrete scale for quantification. When suitable data were collected by the authors but not obtainable from the article, we wrote to the authors and requested the data. If the author cited a dissertation as containing the data, we attempted to locate the data therein because dissertations are not indexed by Web Of Science. We included multivariate trait summaries (e.g., PC axis scores) when reported. If traits reported both the raw trait values and the PC axis scores for a summary of those same traits, we collected both sets of data but omitted the PCs in our main analyses.

Using the above criteria, we screened each article for suitability. As a first pass, we quickly assessed each article for suitability by reading the title and abstract and, if necessary, consulting the main text. After this initial search, we retained 407 studies. Since the previous steps were done by a team of five, a single author (KAT) conducted an in-depth evaluation of each study flagged for possible inclusion. If deemed suitable, we next evaluated whether the necessary data could be obtained. After this second assessment, 198 studies remained. The reasons for exclusion of each study are documented the archived data (see Data accessibility).

### Data collection

For each study, we recorded several types of data. First, we recorded the mean, sample size, and an estimate of uncertainty (if available; e.g., SD or variance) for each measured trait for each parental taxon and hybrid category (cross generation and/or direction). In most cases, these data were included in tables or could be extracted from figures. For figure data extraction, we used ‘WebPlotDigitizer’ (Rohatgi 2019). In some cases, we contacted authors for the raw data or summary data.

Each study contributed a minimum of three records to the larger database: one trait measured in each parent and the F_1_ generation. Traits were categorized as one of: behaviour, chemical, life history, morphological, physiological, or pigmentation. Patterns were generally similar across trait categories and we do not present analyses of the different categories herein. If the same traits were measured over ontogeny, we used only the final data point. When data were reported from multiple ‘trials’ or ‘sites’ we pooled them within and then across sites. If data were reported for different cross directions and/or sexes we recorded data for each cross direction / sex combination separately. We recorded whether each variable was a linear measurement (1D), area measurement (2D), volume measurement (3D; e.g., mass), categorical, or discrete. We did not find meaningful differences in dominance among trait types (ANOVA, *P* = 0.998) and so these data do not factor in to the present analyses. We also recorded the transformation that was applied to the variable, which was used to back-transform traits to the original measurement scale. Data processing was greatly aided by the functions implemented in the `tidyverse` (Wickham 2017).

For each paper we recorded whether the phenotypes were measured in the lab or field, the number of generations of captivity for the parents, and whether a trait correlation matrix (preferably in recombinant—F_2_ or BC_1_—hybrids) was available or calculable from the raw data or figures. We collected trait correlations from every study where they were available, but do not use these data in the present study. Occasionally, different studies analysed different traits of individuals from the same crosses. In these cases, we simply grouped them as being the same cross before analysis.

Calculation of dominance metrics is detailed in the main text. Our main inferences rely on one summary value of each metric per cross. We first calculated dominance for each trait (d_univariate_) or each pair of traits (pairwise d_parent-bias_ and d_mismatch_). We then took the average of these across traits, then across different sexes, and finally across different cross directions.

### Estimating genetic divergence and divergence time

We estimated genetic distance for crosses where sequence data were available for both parents. A preliminary screening revealed that the internal transcribed spacer (ITS I and II) was the most commonly available gene for plants and cytochrome b (cytb) was the most available gene for animals in our dataset. We downloaded sequences in R using the `rentrez` package (Winter 2017), and retained up to 40 sequences per species. Sequences were then aligned with the profile hidden Markov models implemented in the `align` function in the package, `aphid` (Wilkinson 2018). After aligning sequences we calculated genetic distance by simply counting the number of sites that differed between two aligned sequences, implemented using the the `raw` model option in the `dist.dna` function within `ape` (Paradis and Schliep 2018). We made this computation for each pair of sequences and then calculated the average over all pairs for one summary estimate of parental genetic divergence per cross.

We also used timetree (Kumar et al. 2017) to obtain estimates of divergence time for each species pair in their database in years. After obtaining estimates of divergence time we regressed divergence time against the response and predictor variables used in the main analysis. The conclusions from the timetree data do not differ from those using genetic distance and we do not discuss them further (see archived analysis code for these analyses).

### Phylogenetic signal

In the data from the systematic review, we were interested in evaluating whether there was phylogenetic signal in dominance. This might indicate that, for example, dominance was different in plants than in animals. We retrieved NCBI taxonomy IDs for our species using the `taxize R` package (Chamberlain and Szöcs 2013), and used these IDs (one arbitrarily chosen per cross) to generate a phylogeny using phyloT (https://phylot.biobyte.de/). Because branch lengths negligibly affect estimates of phylogenetic signal (Münkemüller et al. 2012), we assigned all branches equal lengths and used the `phylosig` function implemented in `phytools` (Revell 2012) to test for phylogenetic signal via Pagel’s *λ*. Assigning random branch lengths never affected our conclusions. Because we are not testing a causal model, and also because there was no evidence of phylogenetic signal, we do not use the phylogeny in our main text analyses.

### Simulations with no dominance

Estimates of dominance in our analysis could deviate from zero for statistical reasons. For example, any effect of sampling error will bias our estimates of dominance upwards because if the true (scaled) mean of a hybrid is 0.5 then any variation due to sampling error leads to deviation from this value and therefore the appearance of dominance. In addition, biologically real variation around a true mean of zero leads to absolute values greater than zero. We accordingly wished to determine what sorts of values for dominance would be observed if there truly was no dominance.

To determine a null distribution of dominance values if the true dominance were zero, we conducted simulations. For each trait, we generated a simulated vector of normally distributed values with a mean equivalent to the mid-parent value, and sample size and standard deviation equal to the original data. This was done using the back-transformed data in original measurement units. We then took the mean of each vector and used this as the F_1_ mean. Using these data, we regenerated estimates of d_univariate_, and pairwise d_parent-bias_ and d_mismatch_ in the exact same way that was done for the real data. After this, we calculated the median of each dominance metric across all crosses in the simulation. We repeated this process 1000 times to generate 1000 estimates of median dominance values for all three metrics.

The results of these simulations are shown in Fig. S6. These analyses were aided greatly by the `magicfor` R package (Makiyama 2016), which assists with the creation of output objects from `for` loops. By comparing our observed estimates (thick red line in Fig. S6) to the distribution of values without dominance, it is clear that our results largely reflect biological patterns of dominance rather than sampling or statistical artefacts. The simulated dominance values also result from biologically real variation among hybrid individuals due to segregation of heterozygous alleles in the parents and genotype-specific dominance and epistasis.

### Field experiment with sunflowers

Whitney et al. (2006, 2010) generated artificial hybrids to resemble the presumed early ancestors of an existing natural hybrid sunflower, *Helianthus annuus* ssp. *texanus*, which grows in Texas, USA. For further details on the cross, experimental setup and trait measurements, see Whitney et al. (2006, 2010). The BC_1_ generation was obtained by first mating *H. debilis* ssp. *cucumerifolius* from Texas to wild *H. annuus* ssp. *annuus* from Oklahoma to produce F_1_ progeny in the greenhouse. In order to produce enough BC_1_ seed for replicate field populations, a single progeny from the F_1_ generation was propagated vegetatively to produce 14 F_1_ clones. A single *H. a. annuus* pollen donor was mated to the F_1_ clones to produce 3,758 BC_1_ seeds.

To obtain seedlings for the field experiment, seeds were germinated on damp filter paper in late February 2003. Approximately six-day old seedlings were transplanted into peat pots containing field soil and grown in a greenhouse for four weeks before transplanting to the field at the Lady Bird Johnson Wildflower Center, Austin, Texas (hereafter LBJ; 30°10.886’ N, 97°52.58’ W). Prior to planting, plots were tilled to remove standing vegetation. All plants were planted at 90 cm spacing. Plots were fenced with plastic deer fencing to reduce disturbance by deer and rabbits. After planting, local vegetation was allowed to colonise the plots unhindered. Here, we report on 475 BC_1_ individuals planted into a plot (a “selection plot” of Whitney et al. (2006)), as well as individuals of the parental species, *H. a. annuus* (*n* = 44) and *H. debilis* (*n* = 37), planted into a second common-garden plot approximately 500 m away from the BC_1_ plants but within the same site. A photograph of the experiment is included as Fig. S13. A parallel experimenta at a second site, the Brackenridge Field Laboratory, Texas, was analyzed and showed the same results as LBJ, but not included in the present analysis.

Plant traits and fitness were measured from March–September 2 003. Viable seed production was chosen as the measure of fitness in these annual plants. Bags made from plastic mesh (DelStar Technologies, Delaware, USA) were secured onto flowerheads with twist-ties to prevent seed loss in the field. Flowerheads were bagged throughout the season (June to September) to obtain a representative sample. Seed production was estimated by multiplying the total number of heads (bagged & unbagged) by the average number of viable seeds per head in a pooled sample of the bagged heads. In addition to fitness, 30 traits comprising architectural, floral, ecophysiological, phenological, and herbivore resistance traits were measured on each plant, and those traits included herein after filtering are described in Table S 1. Further details on trait measurement protocols and the relevance of individual traits to plant performance are given by Whitney et al. (2006, 2010).

**Table S1.** Table of sunflower traits used to quantify parent-bias and mismatch

| Trait Name | Trait Description and units | Trait Category |
| --- | --- | --- |
| Height of uppermost branch | cm | Architectural |
| Height of lowest branch | cm | Architectural |
| Bushiness | Mean flowerhead position, stem = 1, prim. branch = 2, second. branch = 3 | Architectural |
| Relative Branch Diameter | Mean primary branch basal diameter / stem diameter | Architectural |
| Plant Volume | volume of the main stem (cm<sup>3</sup>) | Architectural |
| Seed weight | Avg mass of individual seed | Architectural |
| Specific leaf area | ratio of leaf area to mass, cm<sup>2</sup> g<sup>-1</sup> | Ecophysiological |
| Water-use efficiency | δ 13C | Ecophysiological |
| Leaf shape | Leaf length : width ratio | Ecophysiological |
| Leaf Carbon:Nitrogen ratio |  | Ecophys. & palat. |
| Disk Diameter | Diameter of the inflorescence disk (mm) | Floral |
| Ligule number |  | Floral |
| Ligule length | cm | Floral |
| Ligule width | cm | Floral |
| Phyllary length | mm | Floral |
| Phyllary width | mm | Floral |
| Bud initiation time | Time between planting and initiation of first inflorescence bud (days) | Phenological |
| Seed maturation time | Period between pollination and seed maturity (days) | Phenological |
| Glandular trichome density | On bottom surface of leaf; these contain sesquiterpene lactones (mm<sup>2</sup>) | Palatability |
| Nonglandular trichome density | On bottom surface of leaf; physical rather than chemical defenses (mm<sup>2</sup>) | Palatability |

## Supplementary material 2: Supplementary figures

**Fig. S1.**
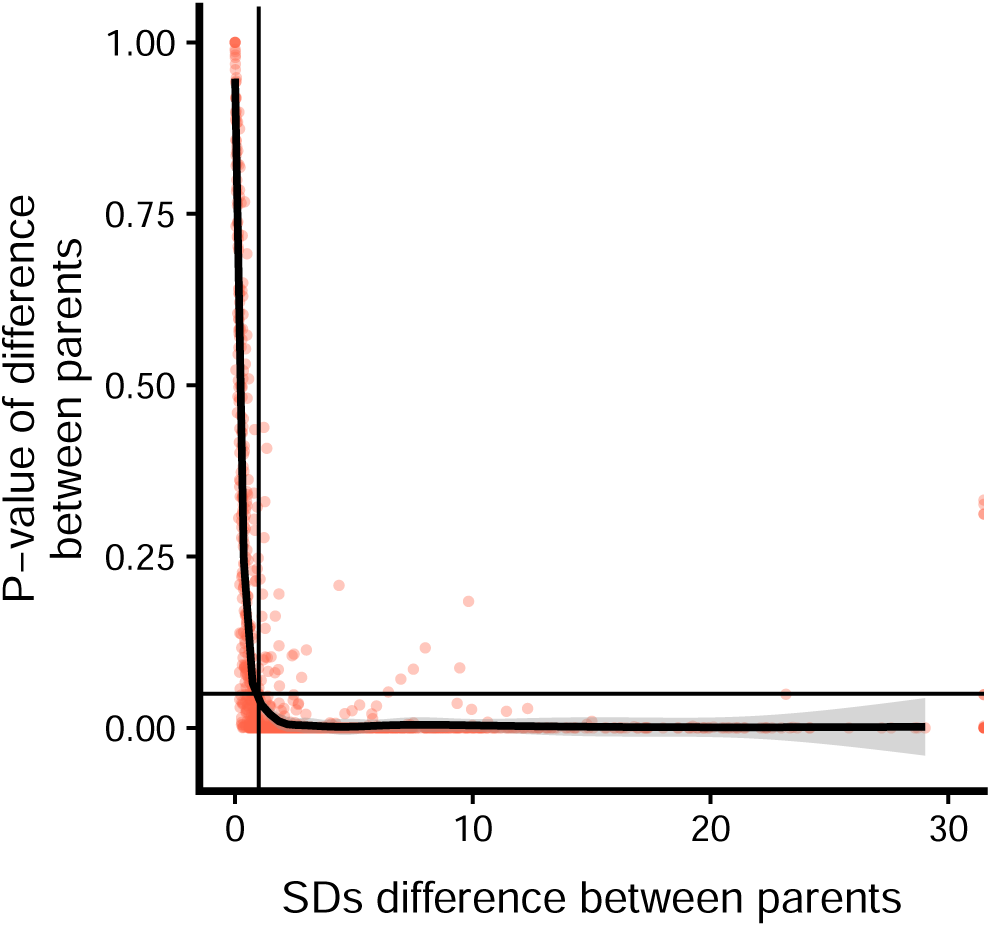
Relationship between SD divergence and *P*-value of *t*-test. Horizontal line is at *P* = 0.05 and the vertical line is at SD = 1. All values except for those in the upper-left quadrant were included in the study. This graph is meant to show that very few traits with > 1 SD divergence have non-significant P-values (upper right quadrat).

**Fig. S2.**
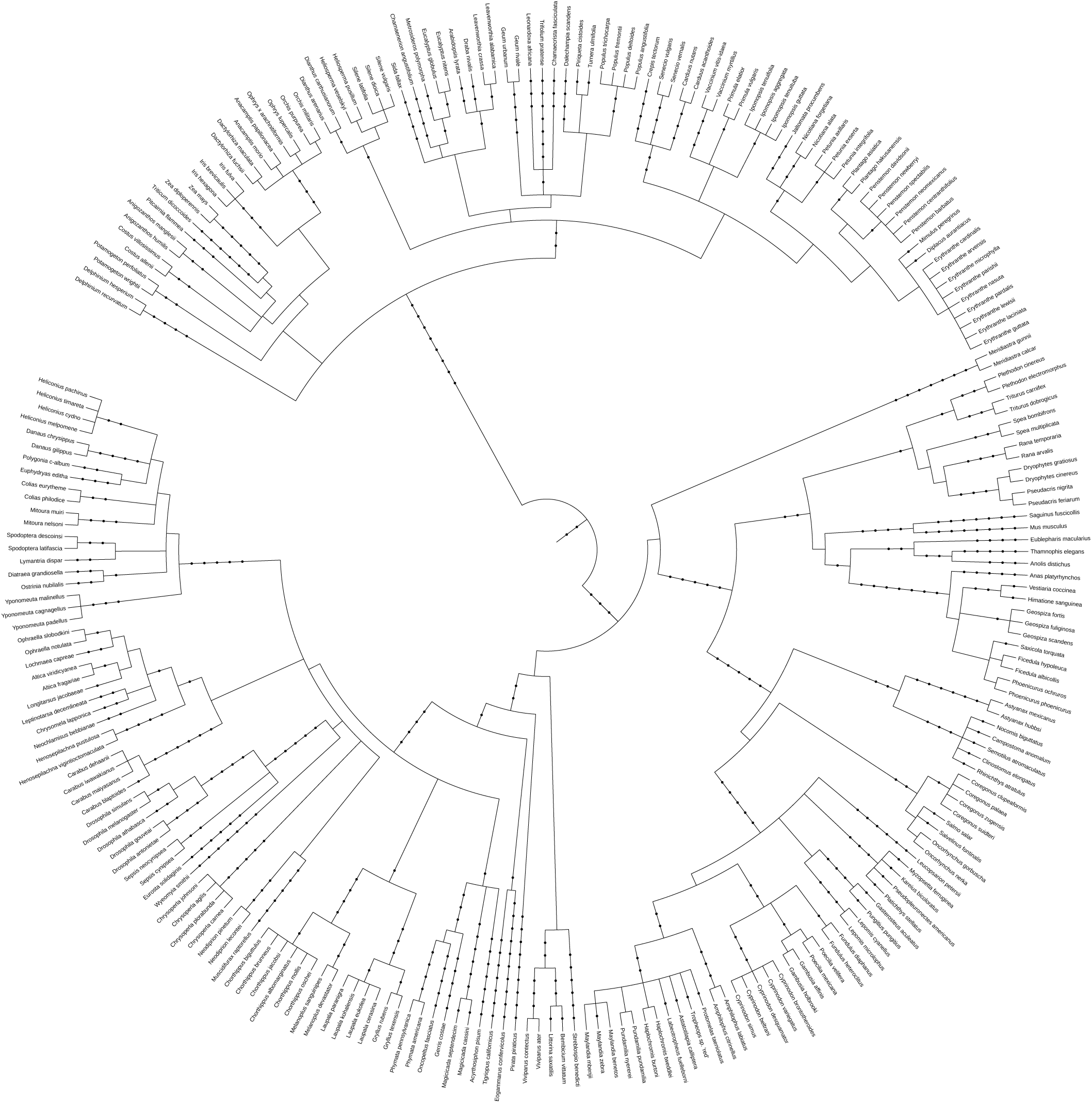
Phylogeny of all species used in this study. For phylogenetic signal analyses, we randomly chose one of the parent species from each pair.

**Fig. S3.**
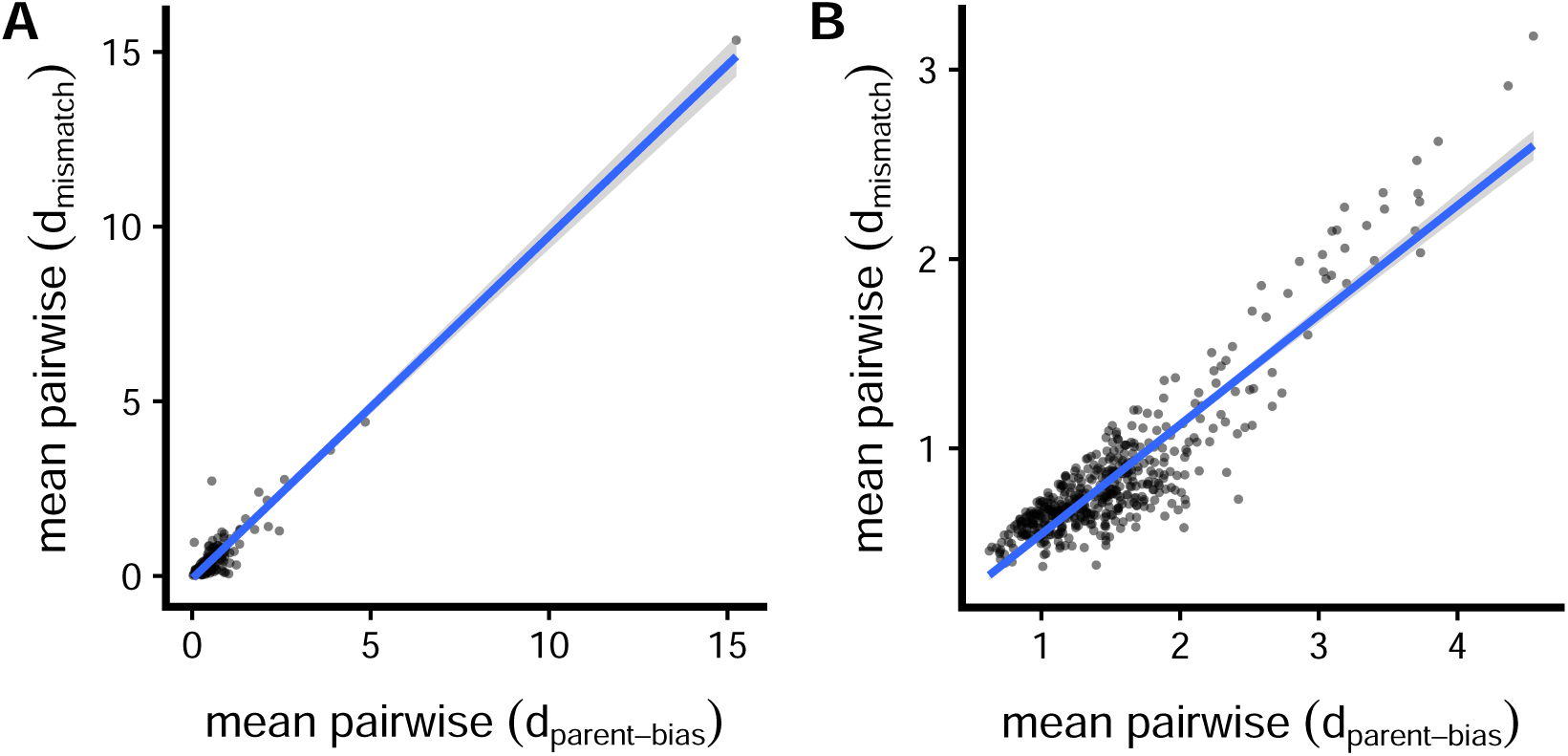
Relationship between mean pairwise parent-bias and mismatch dominance in systematic review and sunflower data. Panel **a** shows the distribution of pairwise d_parent-bias_ and d_mismatch_ in the systematic review data (F_1_s). Panel **B** shows the same metrics calculated at the individual level in the sunflower field experiment data. Both relationships are statistically significant.

**Fig. S4.**
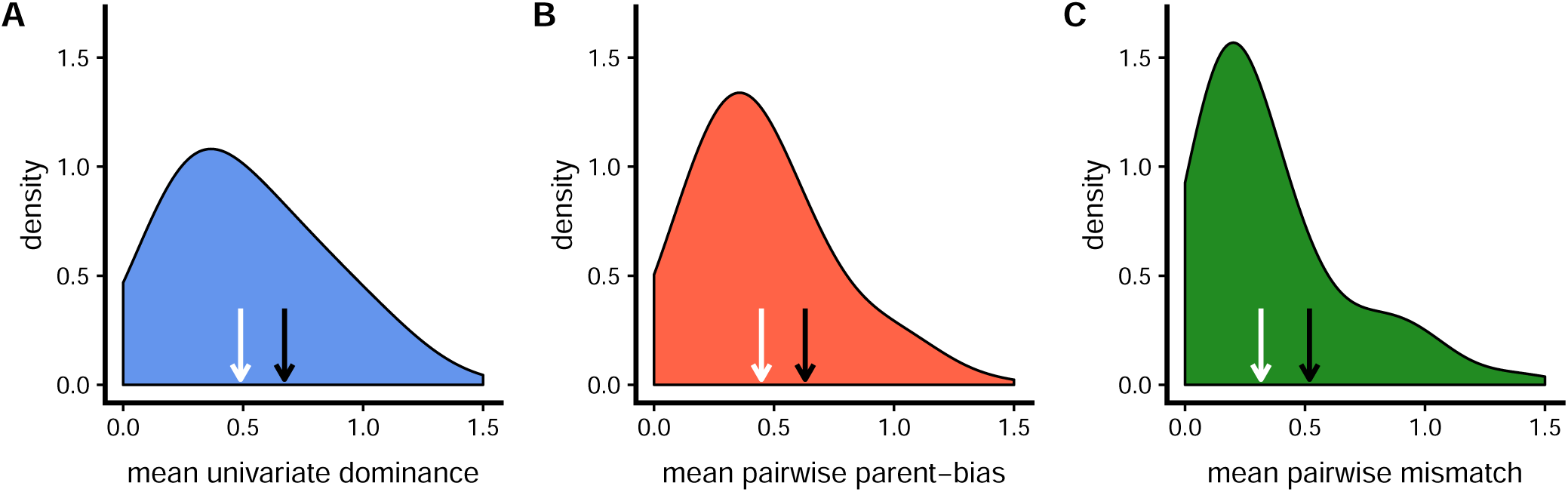
Summary of cross median dominance metrics with each cross contributing a single value. Everything is the same as Fig. 2 of the main text, except here each cross contributed the median value instead of the mean for each dominance metric. The density plots (*y*-axis standardized across panels) show the three main dominance metrics contained herein, with each cross contributing at most a single value per panel. Values of 0 indicate no dominance, values of 1 indicate the maximum without transgression, and values > 1 reflect transgression. The *x*-axis is truncated at 1.5, but the mean of cross medians (black arrows) and median of cross medians (white arrows) are calculated from the whole dataset. Panel **a** shows the univariate dominance (d_univariate_; eqn. 1), panel **b** shows parent-bias (pairwise d_parent-bias_; eqn. 3), and panel **c** shows mismatch (pairwise d_mismatch_; eqn. 4). Panel **a** contains one value from all crosses (*n* = 233) while panels (**b**) and (**c**) only contain information from crosses wherein two or more traits were measured (*n* = 165). Density plots were generated using the `geom_density` function in `ggplot2` with twice the default smoothing bandwidth.

**Fig. S5.**
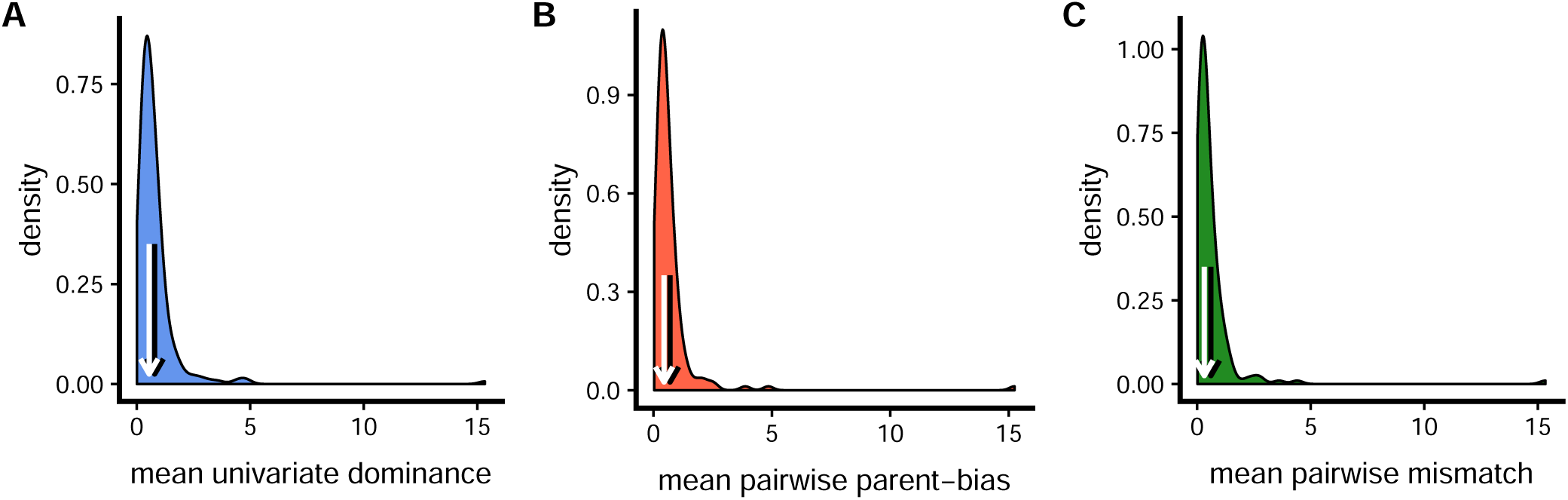
Figure 2 from main text with unconstrained axes. Exactly the same data as in Fig. 2, but without the truncation of *x*-axes at 1.5.

**Fig. S6.**
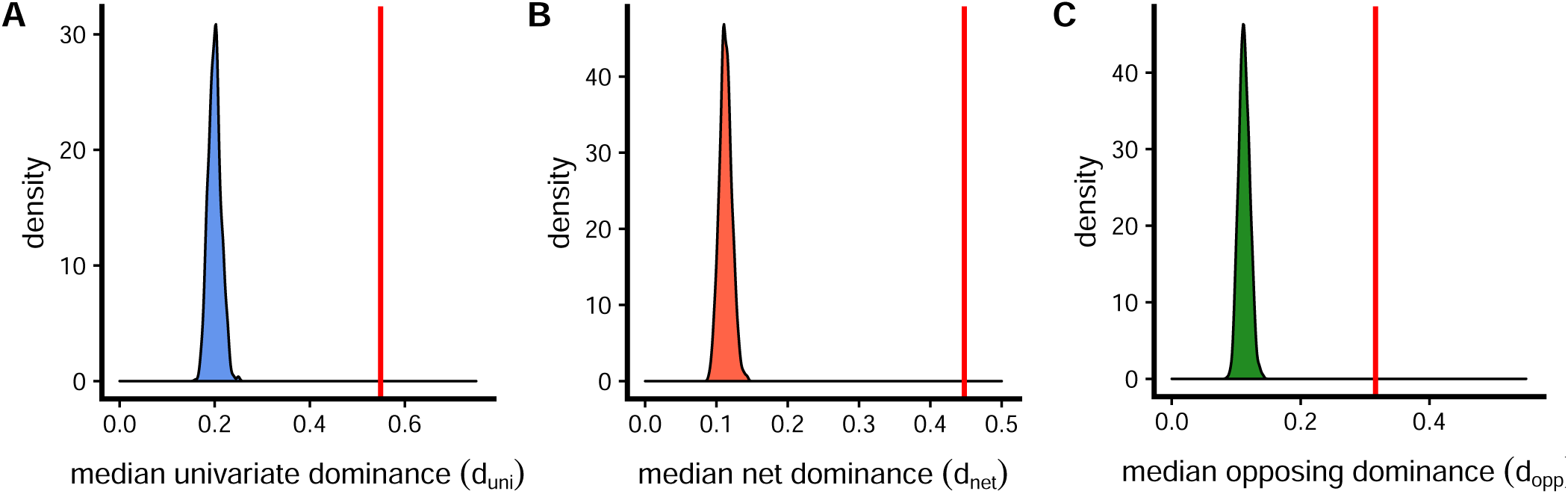
Expected patterns based on sampling error alone (from simulations). Red vertical lines are the median values observed in the main text. We conducted simulations where the true mean was 0 for each trait and individuals were with phenotypes determined by the SD and with identical sample sizes to the real data. The distribution of median estimates, with each simulation (*n* = 1000) contributing one value to each plot are shown as density plots. It is clear that the patterns observed are not just due to sampling error.

**Fig. S7.**
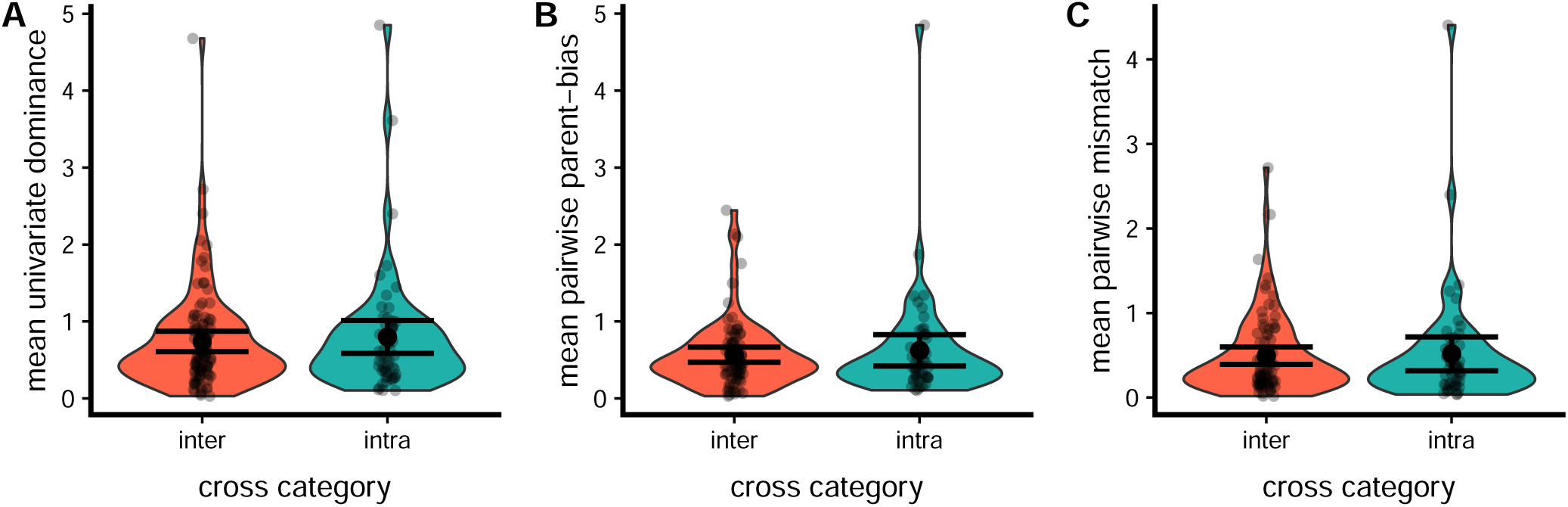
No difference in any dominance metrics between intra-specific and inter-specific crosses across the entire dataset. Each point is the mean dominance metric for d_univariate_ (panel **a**), pairwise d_parent-bias_ (panel **b**), and pairwise d_mismatch_ (panel **c**). All *P* > 0.5.

**Fig. S8.**
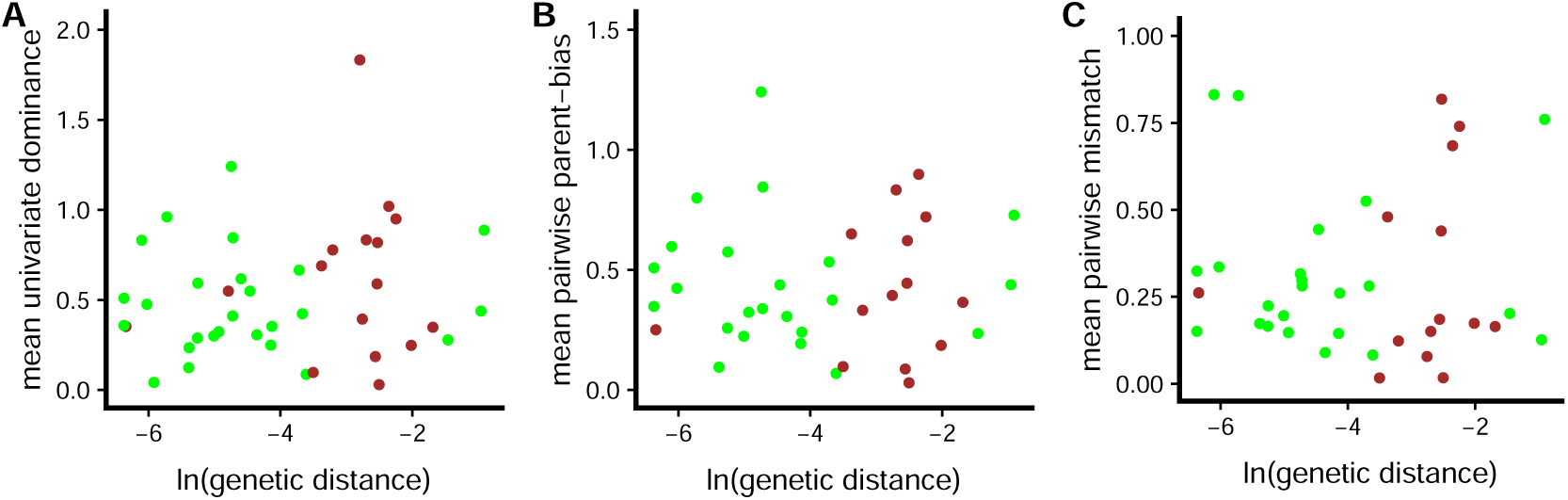
No association between any dominance metrics and genetic distance between the parents. Divergence time was calculated from nucleotide sequences. Each point is the dominance metric for a cross (calculated following eqns 1–4 in the main text). Green points are plants, and brown points are animals. All *P* > 0.5.

**Fig. S9.**
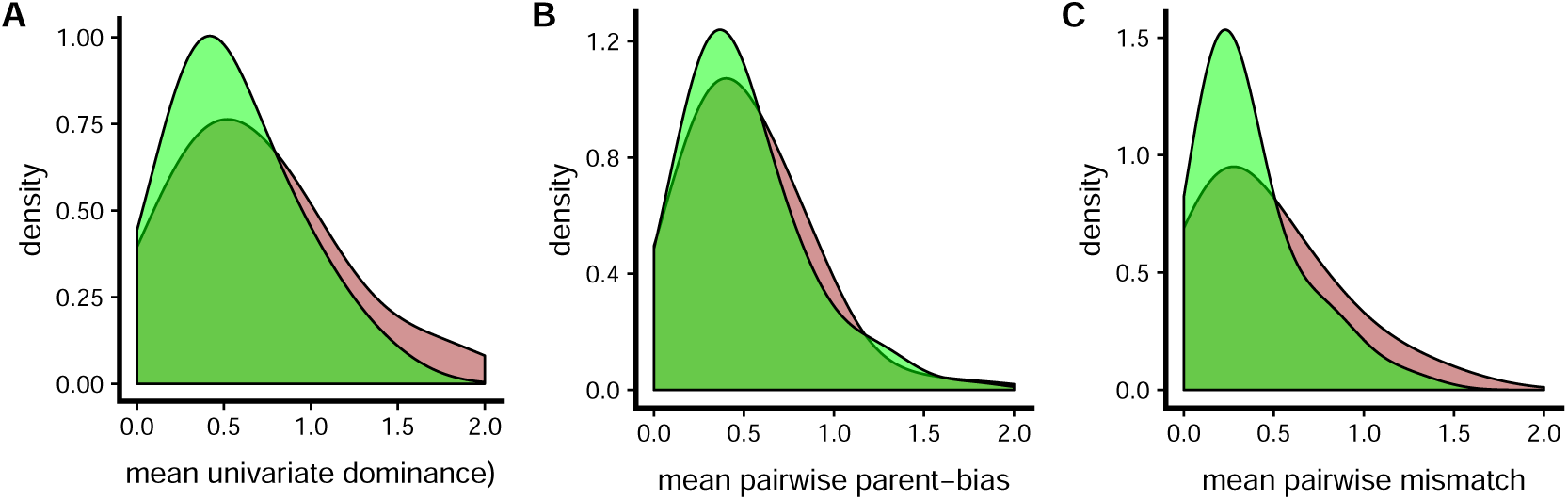
No differences in dominance between plants and animals. Each point is the dominance metric for a cross (calculated following eqns 1–4 in the main text). All *P* > 0.3. Green density plot represents plants, and brown represents animals.

**Fig. S10.**
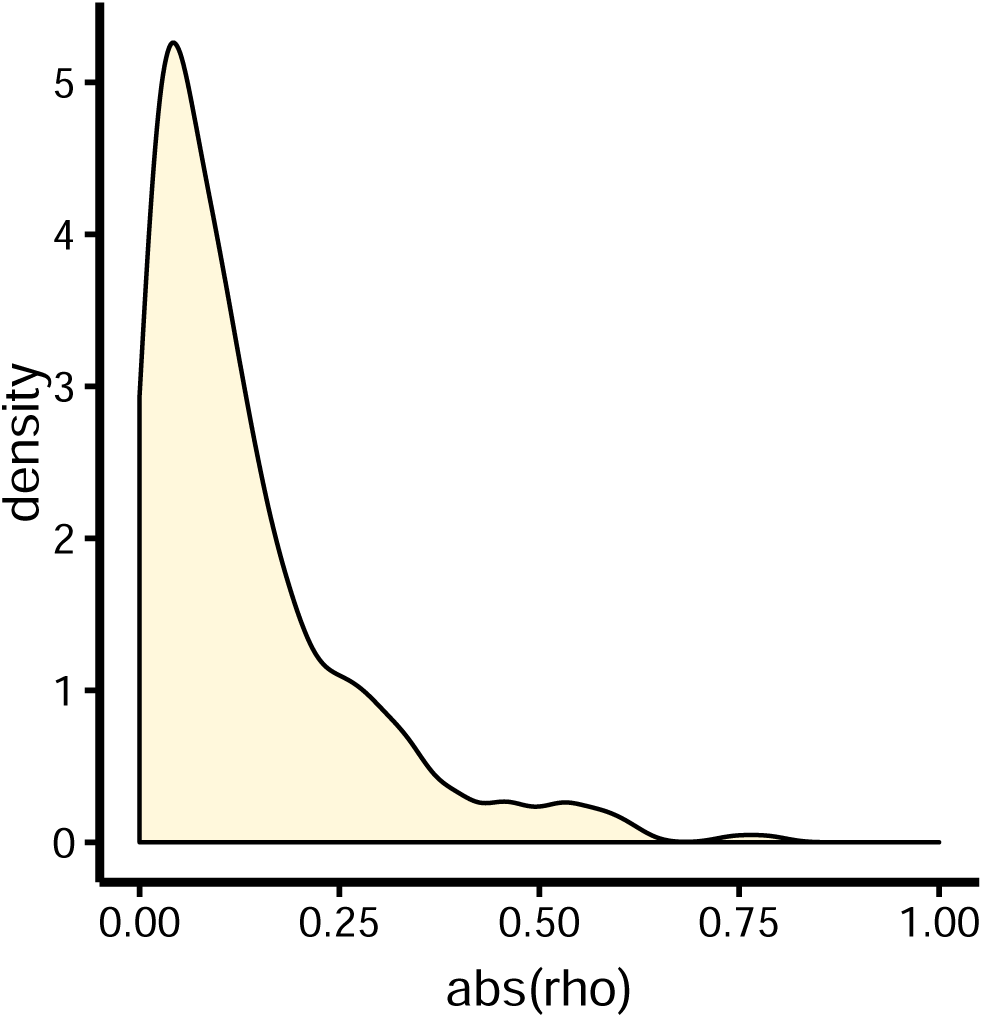
Distribution of pairwise trait correlations in the sunflower data (BC_1_s only). Since most correlations are weak, we conclude that it is unlikely issues caused by trait correlations underlie any of our conclusions.

**Fig. S11.**
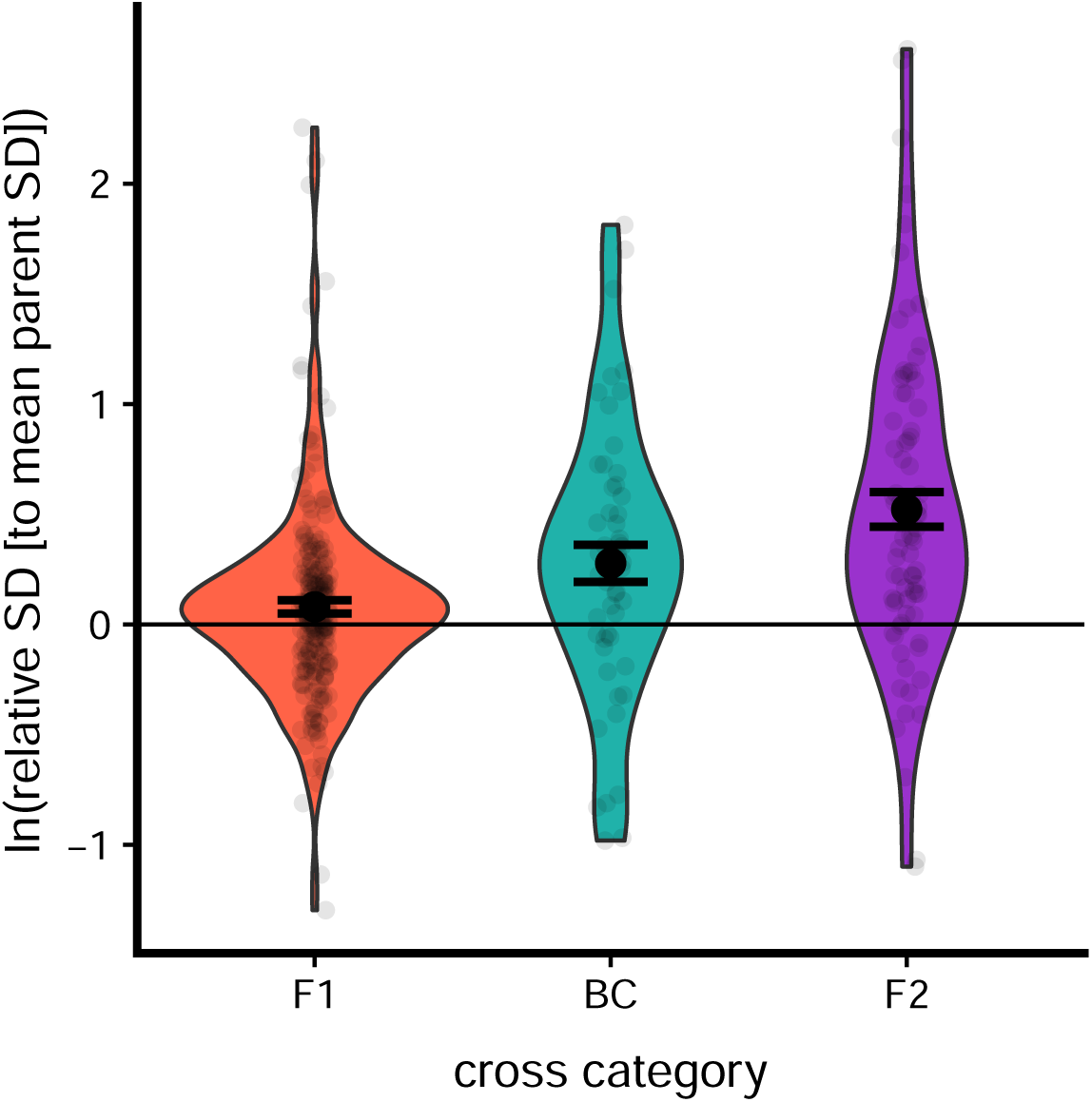
Phenotypic variance in hybrid cross categories relative to mean of parents. The relative SDs are log transformed, so equal variance would be ln(1) = 0. When the difference in phenotypic SD is analyzed in a formal meta-analysis, only the BC_1_ and F_2_ are significantly different compared to parents.

**Fig. S12.**
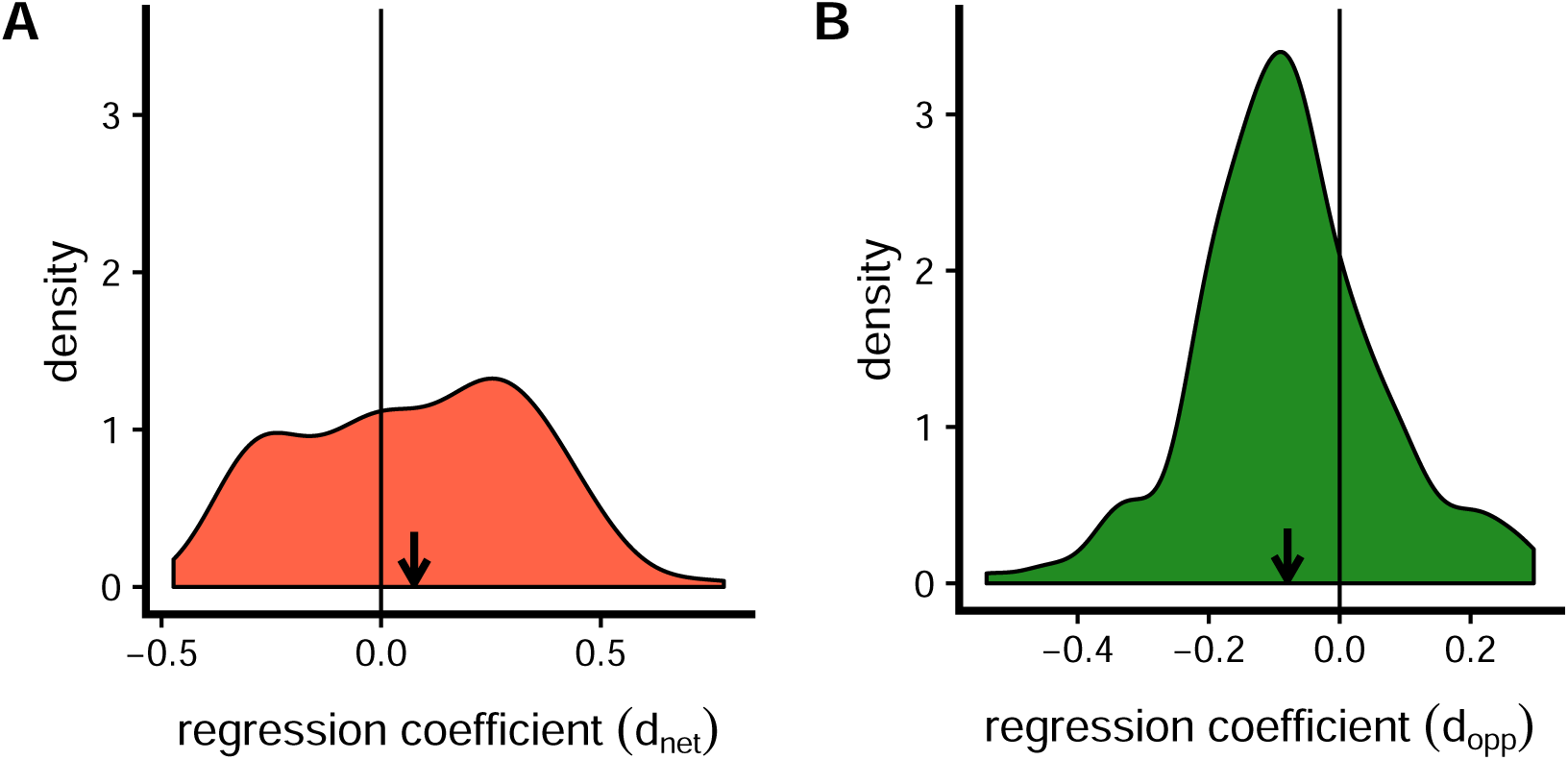
Distribution of regression coefficients in multiple regression analyses (see equation 5). The vertical line demarcates a slope of zero. The black arrows show the mean of pairwise regression coefficients (plotted in the density plot).

**Fig. S13.**
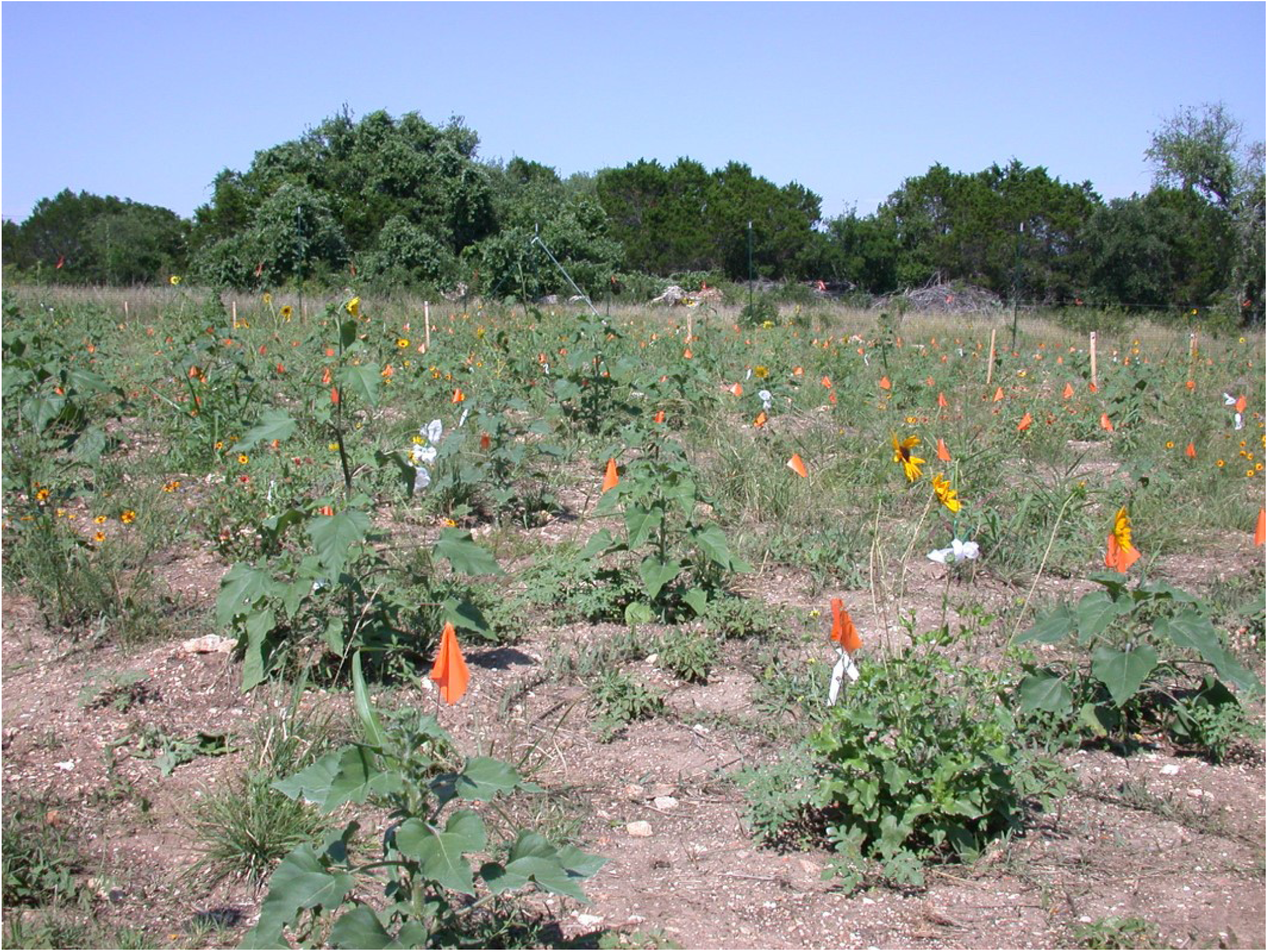
Photograph of sunflower experiment at the Lady Bird Johnson Wildflower Center, in Austin, Texas, USA.

